# COVID-19 genetic risk variants are associated with expression of multiple genes in diverse immune cell types

**DOI:** 10.1101/2020.12.01.407429

**Authors:** Benjamin J. Schmiedel, Vivek Chandra, Job Rocha, Cristian Gonzalez-Colin, Sourya Bhattacharyya, Ariel Madrigal, Christian H. Ottensmeier, Ferhat Ay, Pandurangan Vijayanand

## Abstract

Common genetic polymorphisms associated with severity of COVID-19 illness can be utilized for discovering molecular pathways and cell types driving disease pathogenesis. Here, we assessed the effects of 679 COVID-19-risk variants on gene expression in a wide-range of immune cell types. Severe COVID-19-risk variants were significantly associated with the expression of 11 protein-coding genes, and overlapped with either target gene promoter or *cis*-regulatory regions that interact with target promoters in the cell types where their effects are most prominent. For example, we identified that the association between variants in the 3p21.31 risk locus and the expression of *CCR2* in classical monocytes is likely mediated through an active cis-regulatory region that interacted with *CCR2* promoter specifically in monocytes. The expression of several other genes showed prominent genotype-dependent effects in non-classical monocytes, NK cells, B cells, or specific T cell subtypes, highlighting the potential of COVID-19 genetic risk variants to impact the function of diverse immune cell types and influence severe disease manifestations.

## MAIN

The clinical presentation of SARS-CoV-2 infection in humans can range from severe respiratory failure to disease that is very mild or without symptoms^1^. Although hyperactivation of various cellular components of the immune system have been observed in patients with severe COVID-19 illness^2,3^, the host genetic factors that determine susceptibility to severe COVID-19 illness are not well understood. Genome-wide association studies (GWAS) addressing this question have identified a number of genetic variants associated with COVID-19 susceptibility and severity^4–7^. However, their target genes and the immune cell types where their effects are most prominent are not known. The DICE database of immune cell gene expression, epigenomics and expression quantitative trait loci (eQTLs) (http://dice-database.org) was established to precisely answer these questions as well as to help narrow down functional variants in dense haploblocks linked to disease susceptibility^8,9^. Here, we utilize the DICE database and 3D *cis*-interactome maps to provide a list of target genes and cell types most affected by genetic variants linked to severity of COVID-19 illness.

We systematically assessed the effects of 679 COVID-19-risk variants (defined by the COVID-19 Host Genetics Initiative; release 4 from 20 October 2020^4^; GWAS association *P* value < 5×10^−8^) on gene expression in 13 different immune cell types and 2 activation conditions (**Supplementary Tables 1** and **2**). The expression of 11 protein-coding genes and 1 non-coding RNA (referred here as eGenes) was associated with the genetic variants linked to severe COVID-19 illness requiring hospitalization (**Fig. 1a** and **Extended data Fig. 1**). Notably, the majority of the eGenes associated with severe COVID-19 illness showed prominent effects in specific immune cell types (**Fig. 1b**). Applying a more liberal GWAS association *P* value threshold of 1×10^−5^, we identified 41 additional eGenes that were associated with genetic variants non-significantly linked to severe COVID-19 illness (**Supplementary Table 3**). Some of these variants are likely to reach statistical significance (GWAS association *P* value < 5×10^−8^) as more donors with severe COVID-19 illness are included in the subsequent analysis phases.

**Figure 1.**
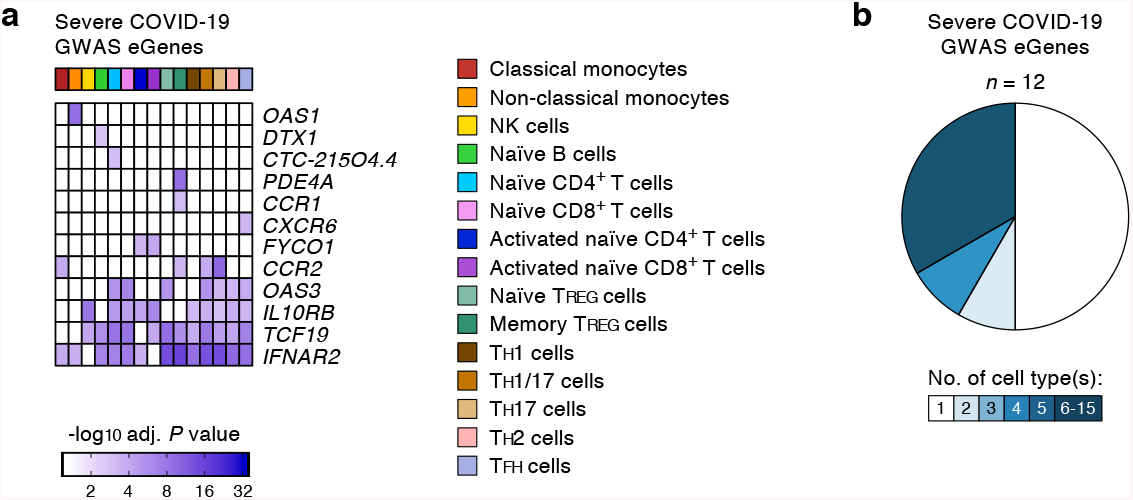
COVID-19-risk variants with eQTL activity in human immune cell types. (**a**) Genes and cell types influenced by GWAS SNPs linked to severe COVID-19 illness requiring hospitalization. For each cell type (columns), the adj. association *P* value for the peak GWAS *cis*-eQTL associated with the indicated eGenes (rows) is shown. (**b**) Fractions of GWAS eGenes linked to severe COVID-19 illness identified in varying numbers of cell types.

Genetic variants in the 3p21.31 locus have been linked to severity of COVID-19 illness by multiple GWAS studies^4–7^. These severe COVID-19-risk variants are inherited as a dense >300 kb haploblock that was shown to have entered the human population >50,000 years ago from Neanderthals^10^. Populations with higher frequency of this Neanderthal-origin COVID-19-risk haplotype have higher risk of severe COVID-19 illness^10^. The severe COVID-19-risk variants in the 3p21.31 locus contains 17 known protein-coding genes (**Fig. 2a**), including *SLC6A20*, *LZTFL1*, *CCR9*, *FYCO1*, *CXCR6*, *XCR1, CCR1*, *CCR3*, *CCR2* and *CCR5*. Among these genes *CCR2* (encoding for C-C type chemokine receptor, also known as monocyte chemoattractant protein 1 receptor) expression showed the strongest association with 3p21.31 severe COVID-19-risk variants identified by GWAS studies^4^ (**Fig. 2a**). Importantly, the risk variants were associated with expression of *CCR2* in certain CD4^+^ memory T cell subsets (T_H_17, T_H_1/17) and classical monocytes (**Fig. 1a**, **2a** and **Supplementary Tables 1** and **2**). Although the *CCR2* promoter did not directly overlap the risk variants, we found that the peak eQTL (rs6808074), located >200kb upstream, directly overlapped an intergenic *cis*-regulatory region that specifically interacted (H3K27ac HiChIP) with *CCR2* promoter in classical monocytes (**Fig. 2b**). These findings suggested that the severe COVID-19-risk variant (rs6808074) likely perturbs the function of a distal enhancer that is important for regulating *CCR2* expression in monocytes. Thus, genetic evidence points to an important role of CCR2 pathway in the pathogenesis of COVID-19. Patients with severe COVID-19 illness were shown to have increased CCR2 expression in circulating monocytes as well as very high levels of CCR2 ligand (CCL2) in bronchoalveolar lavage fluid^11^, supporting the hypothesis that excessive recruitment of CCR2-expressing monocytes may drive pathogenic lung inflammation in COVID-19.

**Figure 2.**
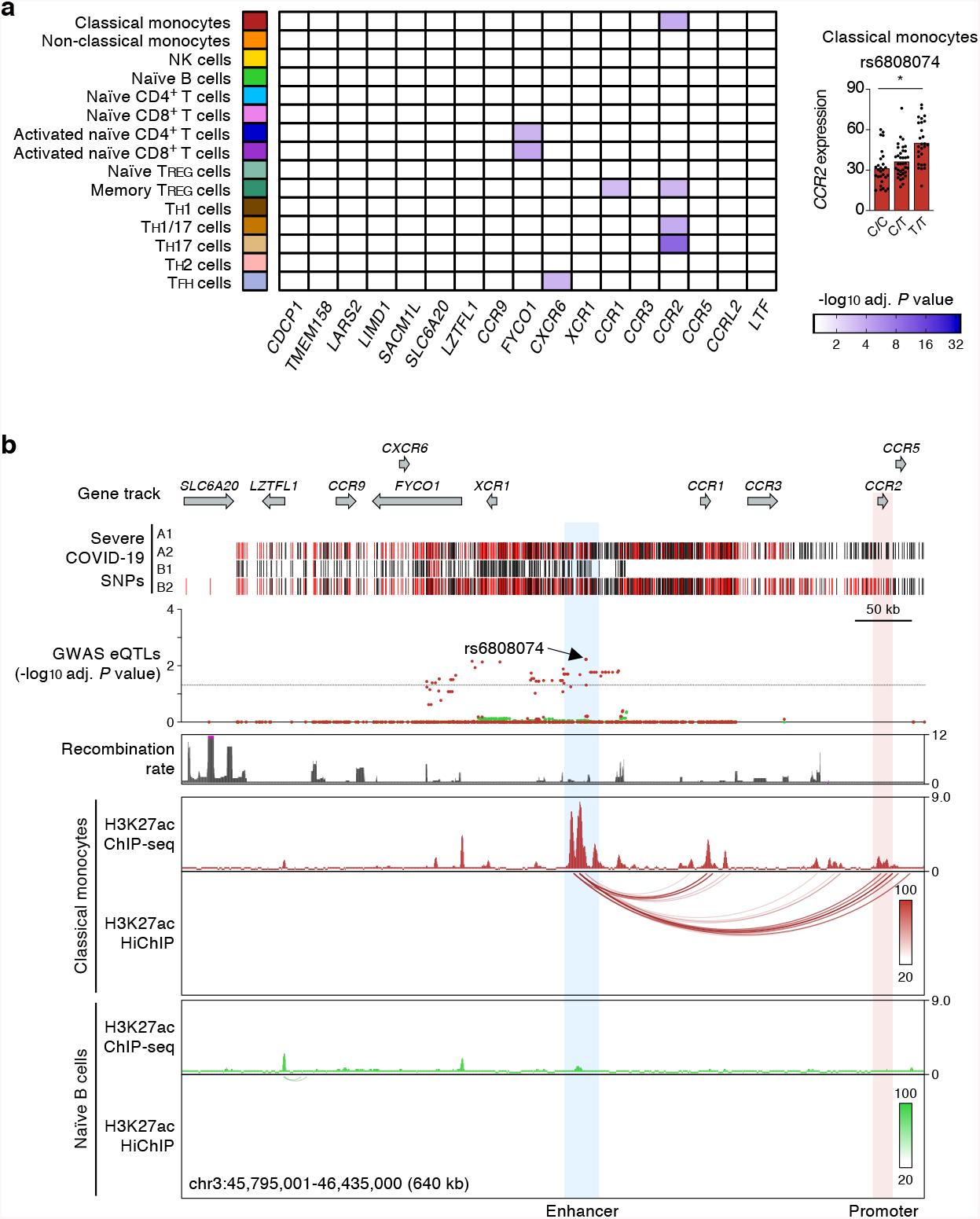
Promoter interacting distal *cis*-eQTLs regulate *CCR2* promoter activity specifically in classical monocytes. (**a**) Genes and cell types most susceptible to the effects of severe COVID-19-risk variants (all with GWAS association *P* value < 5×10^−8^) in the 3p21.31 locus. The adj. association *P* value for the peak GWAS *cis*-eQTL associated with the indicated eGenes in each cell type and activation condition is shown (left). Right, mean expression levels (TPM) of *CCR2* gene in classical monocytes (* adj. association *P* value: 5.94×10^−3^), from subjects (*n*=91) categorized based on the genotype at the indicated GWAS *cis*-eQTL (each symbol represents an individual subject; adj. association P value calculated by Benjamini-Hochberg method). (**b**) WashU Epigenome browser tracks for the 3p21.31 locus, severe COVID-19-risk associated GWAS variants (based on GWAS study, see **Extended Data Figure 1a**; red color bars are lead GWAS SNPs, black color bars are SNPs in linkage disequilibrium), adj. association *P* value for GWAS *cis*-eQTLs associated with expression of *CCR2* expression in classical monocytes (dark red) and naïve B cells (green), recombination rate tracks^27,28^, H3K27ac ChIP-seq tracks, and H3K27ac HiChIP interactions in classical monocytes and naïve B cells.

Defects in the type 1 interferon pathway have been reported in patients with severe COVID-19 illness^12‒15^. We found many severe COVID-19-risk variants in chromosome 21, overlapping the *IFNAR2* gene that encodes for interferon receptor 2, were associated with the expression of *IFNAR2* in multiple immune cell types (**Fig. 3a**). H3K27ac HiChIP-based chromatin interaction maps in this locus showed that the severe COVID-19-risk variants overlapping the *IFNAR2* gene promoter and an intronic enhancer interacted with the promoter of a neighboring gene, *IL10RB* and also influenced its expression levels (**Fig. 3b**). The effects of these risk variants were most prominent in NK cells (rs2284551, adj. association *P* value = 8.99×10^−7^) (**Fig. 3a**). *IL10RB*, encodes for IL-10 receptor beta, and given the immunomodulatory role of IL-10^16^, it is likely that the lower expression on the *IL10RB* in NK cells may perturb their responsiveness to IL-10. Thus, our findings point to a potentially important role for IL-10 signaling and NK cells in influencing susceptibility to severe COVID-19 illness.

**Figure 3.**
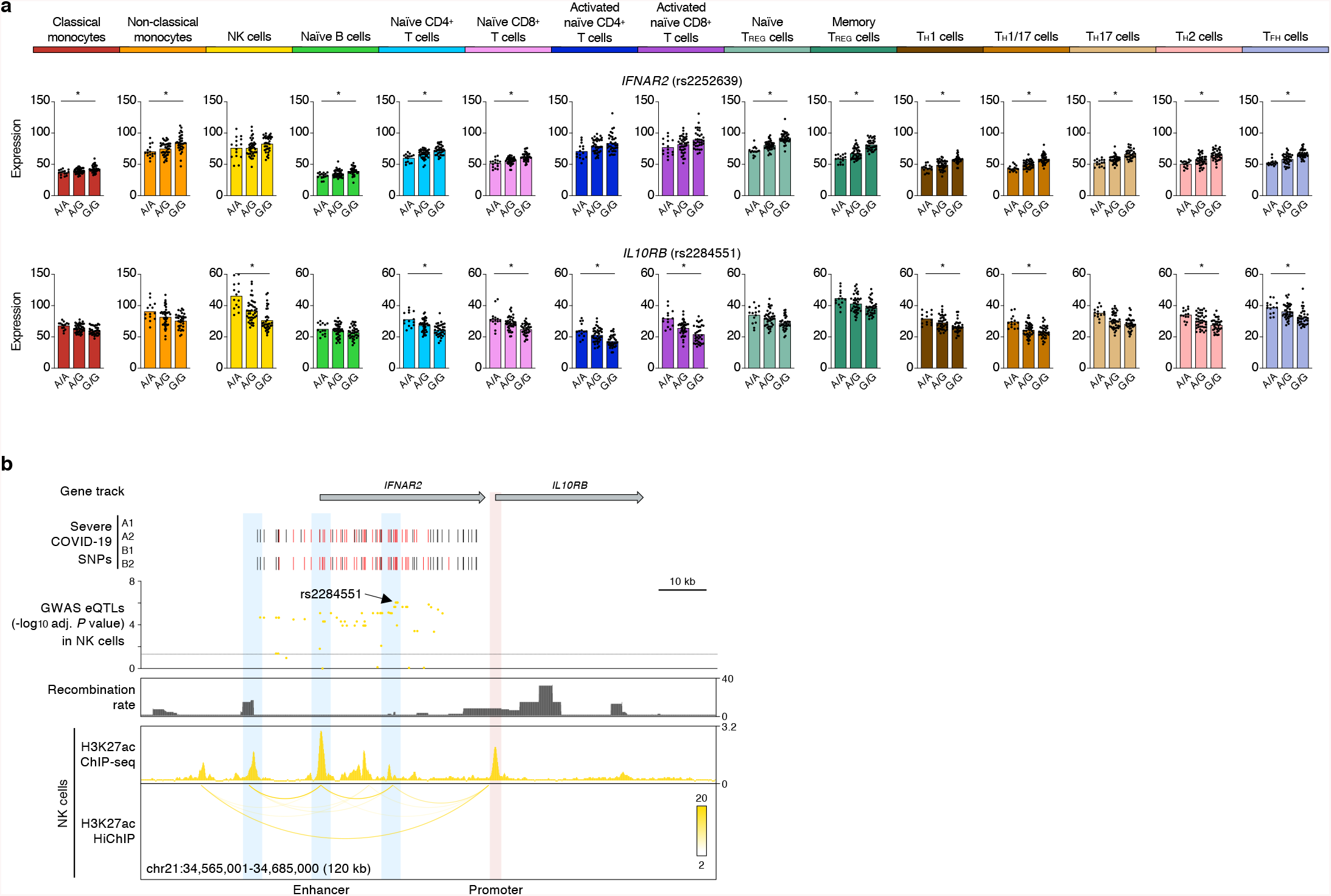
COVID-19-risk variants show cell-type-restriction of their effects on gene expression. (**a**) Mean expression levels (TPM) of selected severe COVID-19-risk associated GWAS eGenes (all with GWAS association *P* value < 5×10^−8^) in the indicated cell types from subjects (*n*=91) categorized based on the genotype at the indicated peak GWAS *cis*-eQTL; each symbol represents an individual subject, * adj. association *P* value < 0.05. (**b**) WashU Epigenome browser tracks for the *IFNAR2* and *IL10RB* loci, severe COVID-19-risk associated GWAS variants (based on GWAS study, see **Extended Data Figure 1a;** red color bars are lead GWAS SNPs, black color bars are SNPs in linkage disequilibrium), adj. association *P* value for GWAS *cis*-eQTLs associated with expression of *IL10RB* in NK cells, recombination rate tracks^27,28^, H3K27ac ChIP-seq tracks, and H3K27ac HiChIP interactions in NK cells.

The expression of two interferon-inducible genes (*OAS1* and *OAS3*) was also influenced by severe COVID-19-risk variants in chromosome 12. *OAS1* and *OAS3*, encode for oligoadenylate synthase family of proteins that degrades viral RNA and activate antiviral responses^17^. *OAS1* showed a peak COVID-19-risk eQTL (rs2057778, adj. association *P* value = 1.77×10^−7^) specifically in patrolling non-classical monocytes, whereas *OAS3* showed prominent eQTLs in T cells (**Fig. 4a**), further highlighting cell-type-restricted effects of severe COVID-19-risk variants. Interestingly, we found that a severe COVID-19-risk variant (rs2010604, adj. association *P* value = 4.50×10^−2^) in the *OAS1*/OAS3 locus also influenced the expression of a neighboring gene (*DTX1*) specifically in naïve B cells (**Fig. 4a**). Active chromatin interaction maps in naïve B cells showed that a *cis*- regulatory region near the eQTL (rs2010604) interacts with the promoter of *DTX1*, located >80kb away, and likely modulates its transcriptional activity (**Fig. 4b**). This notion is supported by recent reports showing that promoters can interact with neighboring gene promoters and regulate their expression^9,18^. *DTX1*, encodes for a ubiquitin ligase Deltex1 that regulates NOTCH activity in B cells^19^. Deltex1 has also been shown to promote anergy, a functionally hypo-responsive state, in T cells^20^; if Deltex1 has a similar functions in B cells, then genetic modulation of *DTX1* levels may have a profound impact on the function of B cells in COVID-19 illness.

**Figure 4.**
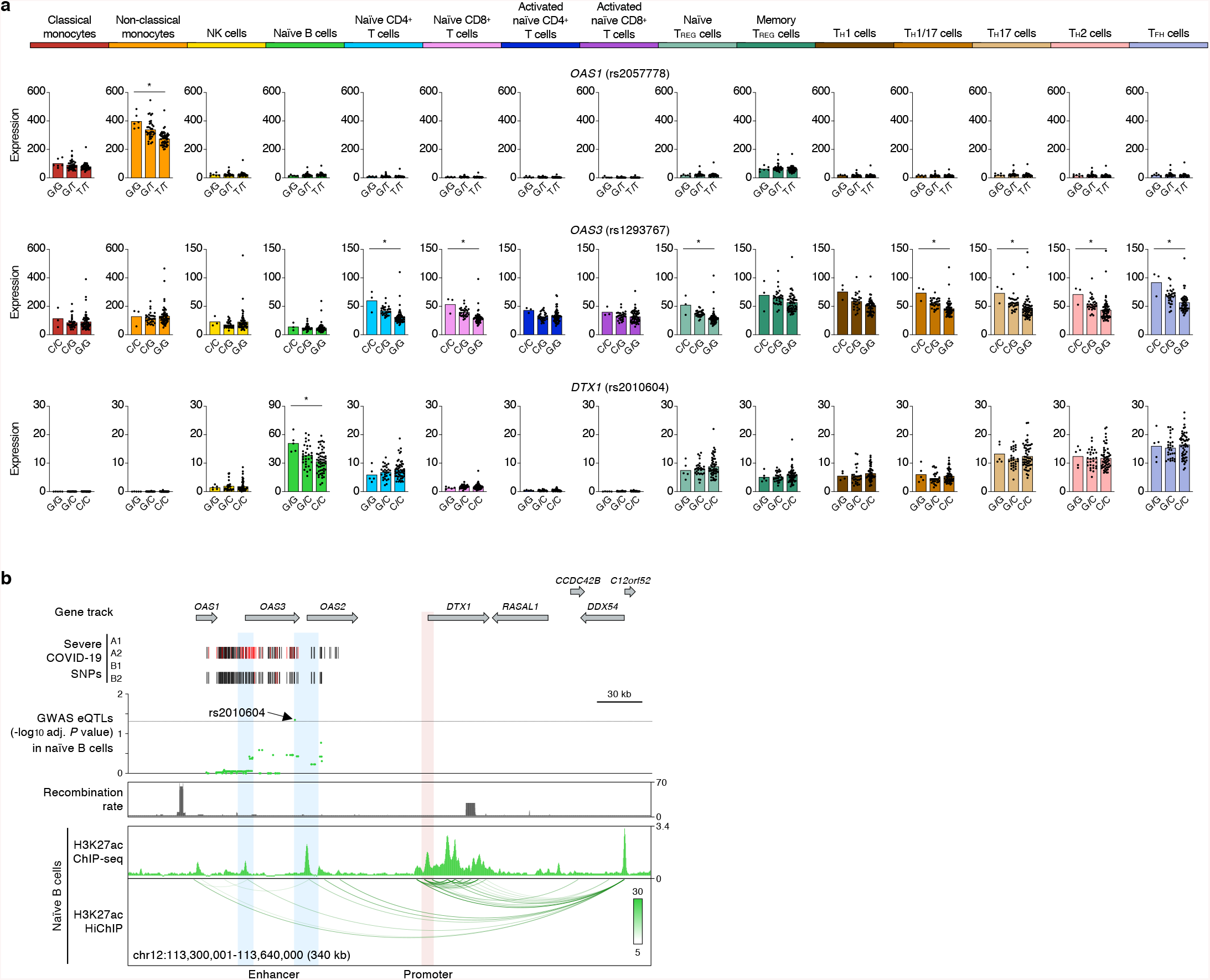
Target genes of severe COVID-19-risk variants in chromosome 12. (a) Mean expression levels (TPM) of selected severe COVID-19-risk associated GWAS eGenes (all with GWAS association *P* value < 5×10^−8^) in the indicated cell types from subjects (*n*=91) categorized based on the genotype at the indicated peak GWAS *cis*-eQTL; each symbol represents an individual subject, * adj. association *P* value < 0.05. (**b**) WashU Epigenome browser tracks for the extended *DTX1* locus, severe COVID-19-risk associated GWAS variants (based on GWAS study, see **Extended Data Figure 1a;** red color bars are lead GWAS SNPs, black color bars are SNPs in linkage disequilibrium), adj. association *P* value for GWAS *cis*-eQTLs associated with expression of *DTX1* in naïve B cells, recombination rate tracks^27,28^, H3K27ac ChIP-seq tracks, and H3K27ac HiChIP interactions in naïve B cells.

Several COVID-19 risk variants located in the promoter region of *TCF19* were associated with its expression in multiple lymphocyte subsets but not in classical or non-classical monocytes (**Fig. 1a**). *TCF19* encodes for a transcription factor TCF19 that has been shown to regulate activation of T cells^21^ and also involved in cell proliferation^22,23^. A noteworthy example of a highly cell-specific severe COVID-19-risk eGene in regulatory T cells (T_REG_) was *PDE4A* (**Extended data Fig. 2**). This gene encodes for phosphodiesterase 4A, which has been shown to reduce the levels of cAMP, and thus influence T cell activity to module immune responses^24^.

In summary, several severe COVID-19-risk variants show cell-type-restriction of their effects on gene expression, and thus have the potential to impact the function of diverse immune cell types and gene pathways. Our analysis of eQTLs and *cis*-interaction maps in multiple immune cell types enabled a precise definition of the cell types and genes that drive genetic susceptibility to severe COVID-19 illness, potentially contributing to the different clinical outcomes. Our study also highlights how information about common genetic polymorphisms can be used to define molecular pathways and cell types that play a role in disease pathogenesis.

## METHODS

The Institutional Review Board (IRB) of the La Jolla Institute for Allergy and Immunology (LJI; IRB protocol no. SGE-121-0714) approved the study. For the DICE study, a total of 91 healthy volunteers were recruited in the San Diego area, who provided leukapheresis samples at the San Diego Blood Bank (SDBB) after written informed consent. Details of gene expression and eQTL analysis in 13 immune cell types and 2 activation conditions have been reported for the DICE project (recalculated to incorporate 4 previously missing RNA-seq samples)^8^, H3K27ac HiChIP data in 5 common immune cell types has been previously reported for the DICE *cis*-interactome project^9^. Genetic variants associated with COVID-19 were downloaded from the COVID-19 Host Genetics Initiative (release 4 from 20 October 2020). Lead genetic variants with GWAS association *P* value < 5×10^−8^ were utilized for downstream analysis. Linkage disequilibrium (LD) for lead COVID-19-risk-variants was calculated using PLINK v1.90b3w^25^ for continental ‘super populations’ (AFR, AMR, EAS, EUR, SAS) based on data from the phase 3 of the 1,000 Genomes Project^26^. SNPs in tight genetic linkage with GWAS lead SNPs (LD threshold R^2^ > 0.8) in any of the five super-populations were retrieved along with the SNP information (*e.g.*, genomic location, allelic variant, allele frequencies). Utilizing this data set, GWAS SNPs (lead SNPs and SNPs in LD) were analyzed for overlap with eQTLs in the DICE database (raw *P* value < 0.0001, adj. association P (FDR) < 0.05, TPM > 1.0) separately for each cell type to identify COVID-19-risk variants that were associated with gene expression in immune cell types.

## DATA AND SOFTWARE AVAILABILITY

The DICE project provides anonymized data for public access at http://dice-database.org. Individual-specific RNA-sequencing, genotype, HiChIP and ChIP-seq data are available from the database of Genotypes and Phenotypes (dbGaP Accession number: phs001703.v3.p1).

## CODE AVAILABILITY

The code used for the analyses performed in this study is available upon request. The codes used for HiChIP data analysis is available on GitHub at https://github.com/ay-lab/pieQTL_NG.

## ACKNOWLEDGMENTS

This work was funded by NIH grants R24-AI108564 (P.V., F.A., C.H.O.), the William K. Bowes Jr Foundation (P.V.), and R35-GM128938 (F.A.).

## AUTHOR CONTRIBUTIONS

B.J.S., V.C., C.H.O., F.A., and P.V. conceived the work. B.J.S, V.C., F.A., and P.V. designed the study and wrote the manuscript. J.R., C.G.-C., S.B., and A.M., performed bioinformatic analyses under the supervision of B.J.S., V.C., P.V. and F.A.

## EXTENDED DATA FIGURE LEGENDS

**Extended data Figure 1.**
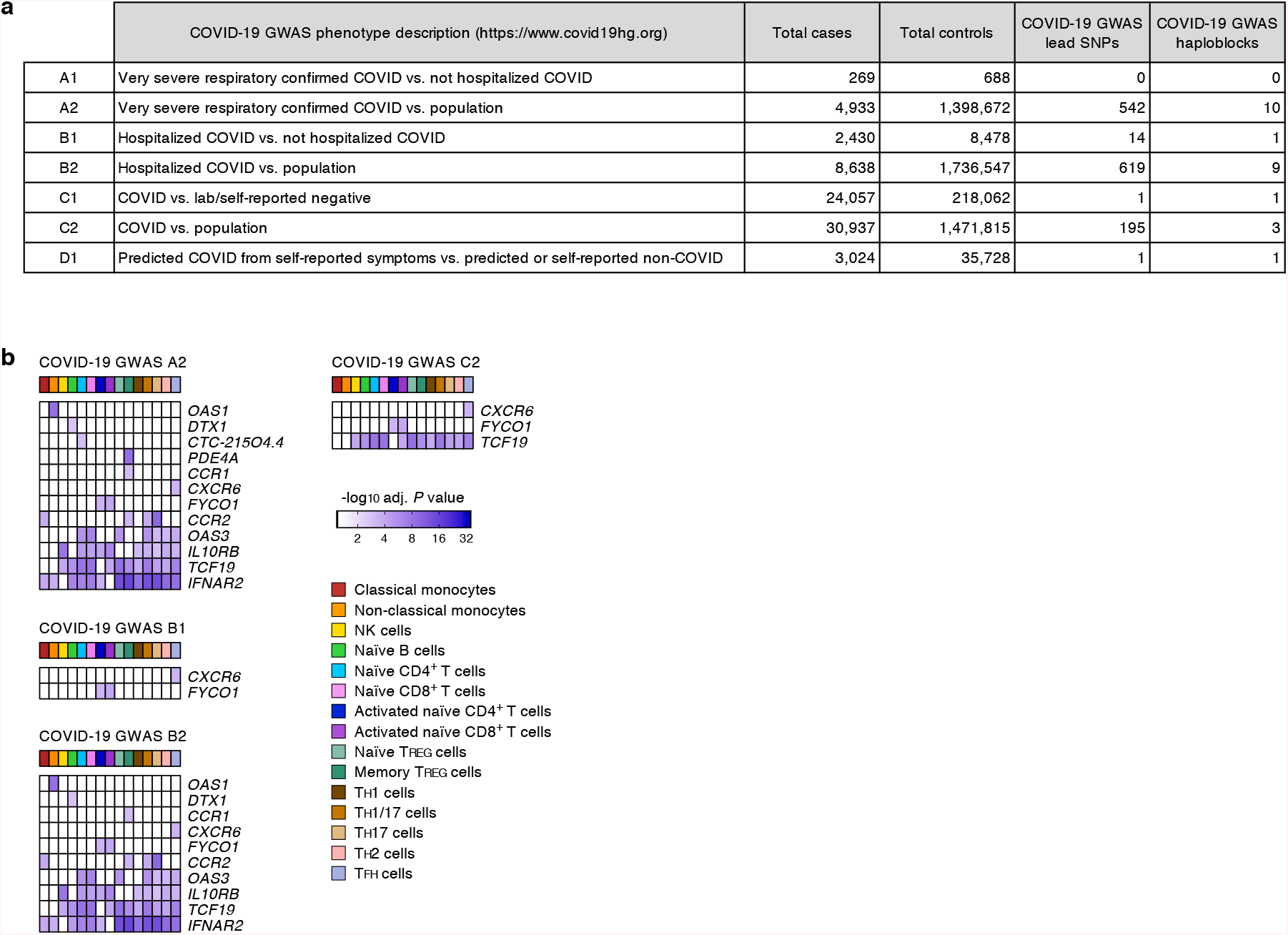
Gene and cell types most susceptible to severe COVID-19-risk associated GWAS SNPs. (**a**) GWAS SNP datasets defined by the COVID-19 Host Genetics Initiative (see **Online Methods**), number of cases and controls in each study (release 4 from 20 October 2020), retrieved GWAS lead SNPs (GWAS association *P* value < 5×10^−8^) and number of GWAS haploblocks. (**b**) For each separate GWAS SNP dataset (as defined by the COVID-19 Host Genetics Initiative), the adj. association *P* value for the peak GWAS *cis*-eQTL associated with the indicated eGenes in each cell type and activation condition is shown.

**Extended data Figure 2.**
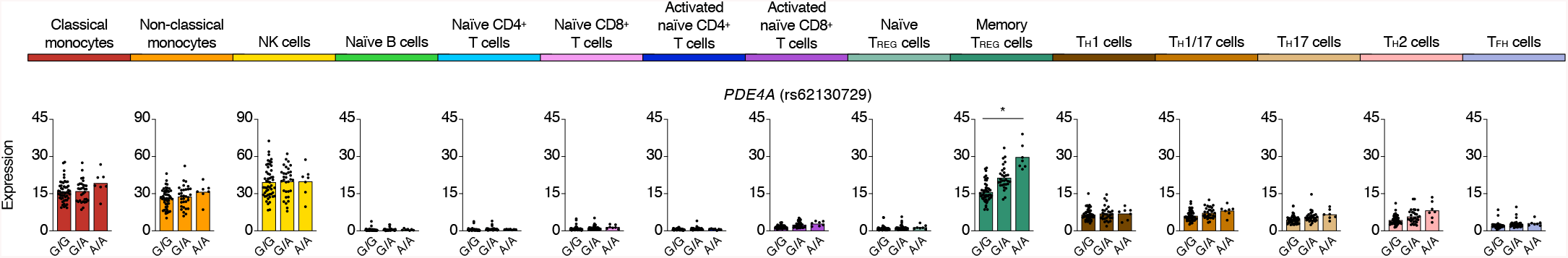
Genes and cell types most susceptible to severe COVID-19-risk associated GWAS variants. Mean expression levels (TPM) of *PDE4A,* a severe COVID-19-risk associated GWAS eGene (GWAS association *P* value < 5×10^−8^), in the indicated cell types from subjects (*n*=91) categorized based on the genotype at the indicated peak GWAS *cis*-eQTL; each symbol represents an individual subject, * adj. association *P* value < 0.05.

## SUPPLEMENTARY TABLES

**Table S1.** List of eGenes in each immune cell type and activation condition, along with information on its peak GWAS *cis*-eQTL (GWAS association *P* value < 5×10^−8^) that is associated with COVID-19 illness.

**Table S2.** List of eGenes in each immune cell type and activation condition, along with information on all GWAS *cis*-eQTLs (GWAS association *P* value < 5×10^−8^) that are associated with COVID-19 illness.

**Table S3.** List of eGenes in each immune cell type and activation condition, along with information on all GWAS *cis*-eQTLs (GWAS association *P* value < 1×10^−5^) that are associated with COVID-19 illness.

